# High Affinity Chimeric Antigen Receptor with Cross-Reactive scFv to Clinically Relevant EGFR Oncogenic Isoforms

**DOI:** 10.1101/2021.02.04.429797

**Authors:** Radhika Thokala, Zev A. Binder, Yibo Yin, Logan Zhang, Jiasi Vicky Zhang, Daniel Y. Zhang, Michael C. Milone, Guo-li Ming, Hongjun Song, Donald M. O’Rourke

## Abstract

Tumor heterogeneity is a key reason for therapeutic failure and tumor recurrence in glioblastoma (GBM). Our chimeric antigen receptor (CAR) T cell (2173 CAR T cells) clinical trial (NCT02209376) against Epidermal growth factor receptor (EGFR) variant III (EGFRvIII) demonstrated successful trafficking of T cells across the blood brain barrier into GBM active tumor sites. However, CAR T cell infiltration was associated only with a selective loss of EGFRvIII+ tumor, demonstrating little to no effect on EGFRvIII^-^ tumor cells. Post-CAR T treated tumor specimens showed continued presence of EGFR amplification and oncogenic EGFR extracellular domain (ECD) missense mutations, despite loss of EGFRvIII. To address tumor escape, we generated an EGFR-specific CAR by fusing monoclonal antibody (mAb) 806 to a 4-1BB co-stimulatory domain. The resulting construct was compared to 2173 CAR T cells in GBM, using in vitro and in vivo models. 806 CAR T cells specifically lysed tumor cells and secreted cytokines in response to amplified EGFR, EGFRvIII, and EGFR-ECD mutations in U87MG cells, GBM neurosphere-derived cell lines, and patient-derived GBM organoids. 806 CAR T cells did not lyse fetal brain astrocytes or primary keratinocytes to a significant degree. They also exhibited superior antitumor activity in vivo when compared to 2173 CAR T cells. The broad specificity of 806 CAR T cells to EGFR alterations gives us the potential to target multiple clones within a tumor and reduce opportunities for tumor escape via antigen loss.

## Introduction

Chimeric antigen receptor (CAR) cells targeting pediatric B cell malignancies have shown unprecedented responses and were the first CAR T cell therapies to receive FDA approval, in 2017 (1–3). The successful application of this therapeutic technology in the treatment of solid tumors, including glioblastoma (GBM), remains a significant challenge; chief among them are tumor heterogeneity, immunosuppressive tumor microenvironment, and antigen escape (4, 5). Successful strategies for overcoming these obstacles are required to advance CAR T therapy in solid tumors.

Epidermal growth factor receptor (EGFR) was one of the first oncogenes identified in GBM and presents an attractive therapeutic target, given its extracellular nature and frequent alterations in GBM. Approximately 60% of GBM specimens contain a mutation, rearrangement, splicing alteration, and/or amplification of EGFR (6). EGFR overexpression, mediated through focal amplification of the EGFR locus as double minute chromosomes, has long been recognized as the most common EGFR alteration, present in 60% of GBM patients (7, 8). Tumor-specific EGFR variant III (EGFRvIII), resulting from deletion of exon 2-7 of wildtype EGFR (wtEGFR), is present in 30% of GBM patients (9). In addition, oncogenic missense mutations EGFR^A289D/T/V^, EGFR^R108G/K^, and EGFR^G598V^ have been identified in 12-13% of cases in the extracellular domain (ECD) of EGFR, independent of EGFRvIII. Missense mutations and EGFRvIII often co-occur with EGFR amplification and activate EGFR receptor independent of its ligand (10). Several of the missense mutations have been shown to have a negative effect on patient survival, driving tumor proliferation and invasion (11).

Our first-in-man CAR T clinical trial (NCT02209376) against EGFRvIII in recurrent GBM demonstrated the safety of a peripheral infusion of CAR T cells and resulted in successful trafficking of the CAR T cells to active tumor sites, across the blood brain barrier (12). After treatment, CAR T cells infiltrated the GBM tumors rapidly, proliferated in situ, and persisted over a prolonged period of time. However, CAR T cell infiltration was associated only with a selective loss of EGFRvIII+ GBM cells. Importantly, post-CAR T treated tumor specimens showed the continued presence of EGFR amplification and missense mutations, despite the decrease in EGFRvIII target antigen. Persistence of EGFR amplification and ECD missense mutations in the context of loss of EGFRvIII expression suggested tumor heterogeneity played an essential role for tumor recurrence and continued regrowth.

mAb806, originally raised against EGFRvIII, recognizes a conformationally exposed epitope of wtEGFR when it is overexpressed on tumor cells. The same epitope is not exposed in EGFR expressed on normal non-overexpressing cells (13, 14). ABT-414, an antibody-drug conjugate composed of a humanized mAb806 (ABT-806), showed early efficacy in phase I/II clinical trials with no apparent skin toxicity in treated GBM patients (15). However, a recent Phase III trial was terminated when an interim analysis failed to demonstrate a survival benefit over placebo (16). mAb806 showed an increased binding affinity for not only EGFRvIII but also EGFR ECD mutations and a low affinity for wtEGFR (11). These findings suggest that mAb806 is a viable therapeutic option for tumors harboring EGFR alterations in addition to EGFRvIII.

In the present study, we have developed EGFR-specific CAR T cells derived from the single chain fragment variable region (scFv) of 806 mAb, using our standard 4-1BB-ζ construct (17). We then compared 806 CAR T activity with EGFRvIII-specific CAR T cells (2173 CAR T), currently in clinic, for specificity against oncogenic EGFR alterations, including amplified EGFR, EGFRvIII, and extracellular mutations *in vitro* and *in vivo*.

## Materials and Methods

### CAR constructs

806 scFvs were swapped with scFv of our standard CD19-BB-ζ lentiviral vector described previously to generate 806-BB-ζ CAR (17, 18). Briefly, the nucleotide coding sequences of 806 or C225 scFv with huCD8 leader were synthesized by Geneart (Thermo Fisher Scientific, Waltham, MA) with 5’ Xba1 and 3’Nhe1 and ligated to Xba1 and Nhe1 sites of CD19-BB-ζ car construct. The C225-BB-ζ CAR was obtained from Dr. Avery Posey’s lab at the University of Pennsylvania. The 2173-BB-ζ CAR T construct was obtained from Dr. Laura Johnson’s Lab at the University of Pennsylvania (19, 20).

### Transduction and expansion of primary human T lymphocytes

Human primary total T cells (CD4 and CD8) were isolated from normal healthy donors following leukapheresis by negative selection using RosetteSep kits (Stemcell Technologies, Vancouver, CA). All specimens were collected with protocol approved by University Review Board and written informed consent was obtained from each donor. T cells were cultured in RPMI 1640 (Thermo Fisher Scientific) supplemented with 10% fetal bovine serum (FBS) (VWR, Radnor, PA), 10 mM HEPES (Thermo Fisher Scientific), 100 U/mL penicillin (Thermo Fisher Scientific), 100 g/mL streptomycin sulfate (Thermo Fisher Scientific), and stimulated with magnetic beads coated with anti-CD3/anti-CD28 (Thermo Fisher Scientific) at 1:3 T cell to bead ratio. Approximately 24 hours after activation, T cells were transduced with lentiviral vectors encoding CAR transgene at an MOI of 3 to 6. On day 5 beads were removed and thereafter cells were counted and fed every 2 days, supplemented with IL 2 150 U/mL until they were either used for functional assays or cryopreserved for future use.

### Cell lines and cell culture

The human cell line U87MG was purchased from the American Type Culture Collection (ATCC) and maintained in MEM (Richter’s modification) (Thermo Fisher Scientific) with components GlutaMAX-1 (Thermo Fisher Scientific), HEPES pyruvate, and penicillin/streptomycin supplemented with 10% FBS. Primary human keratinocytes were purchased from the Dermatology Core Facility at the University of Pennsylvania. K562 cells were purchased from ATCC and maintained in RPMI media (Invitrogen, Carlsbad, CA) supplemented with 10% FBS, 20 mM HEPES, and 1% penicillin/streptomycin. Primary astrocytes were purchased (Sciencell Research Laboratories, Carlsbad, CA) and cultured according to manufacture instructions. The cells from early passages were used for cytotoxicity and cytokine experiments. GSC cell lines were cultured in DMEM F12 media (Sigma Aldrich, St. Louis, MO) supplemented with 2% B27 without vitamin A (Thermo Fisher Scientific), 20 mM HEPES, and penicillin/streptomycin.

### EGFR-mutant cell lines

To produce the overexpressing EGFR cell line (designated as U87MG-EGFR), lentivirus co-expressing wtEGFR and Cyan Fluorescent Protein (CFP) under the control of EF-1α promoter was transduced into U87MG cell line. On post-transduction day 4, cells were sorted on an Influx cell sorter (BD, Franklin Lakes, NJ) on the basis of high EGFR expression, and subsequently expanded. Lentivirus co-expressing CFP and EGFR mutants EGFR^R108K/G^, EGFR^A289D/T/V^, or EGFRvIII were transduced into U87MG-EGFR, GSC5077 neurosphere cells (21), and K562 cell lines. CFP positive cells were sorted by fluorescence activated cell sorting (FACS). For luciferase killing assays and in vivo tracking studies, U87MG and U87MG-EGFR mutant cell lines were transduced with lentivirus click beetle green (CBG) luciferase and Green Fluorescent protein (GFP). Anti-GFP positive cells were sorted by FACS.

### Cytokine analysis

CAR T cells and K562 targets expressing EGFR and its variants were co-cultured in 1:2 ratio in R10 medium in a 96 well plate, in triplicate. Plates were incubated at 37°C with 5% CO_2_. After 48 hours, supernatants were collected and cytokine levels were assessed by ELISA kit (R&D Systems, Minneapolis, MN) for IFN-γ, TNF-α, and IL2 production, according to manufacture instructions.

### Chromium release assay

The cytolytic efficacy of CAR T cells against K562 cells was evaluated by 4-hour chromium release assays using E:T ratios of 5:1, 2.5:1, and 1:1. 51Cr labeled target cells were incubated with CAR T cells in complete medium or 0.1% Triton X-100, to determine spontaneous and maximum 51Cr release respectively, in a V-bottomed 96-well plate. The mean percentage of specific cytolysis of triplicate wells was calculated from the release of ^51^Cr using a Top Count NXT (Perkin-Elmer Life and Analytic al Sciences, Inc., Waltham, MA) as:

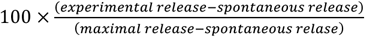

Data was reported as mean ± SD.

### Luciferase based Cytotoxic Assay

CBG+ target cell lines (U87 variants and GSC5077 variants) were co-cultured with CAR T cells at E:T ratios of 10:1, 5:1, and 2.5:1, for 24 hours at 37°C. 100 μL of the mixture was transferred to a 96 well black luminometer plate, 100uL of 66 μg/mL D-luciferin (Goldbio, St. Louis, MO) was added, and the luminescence was immediately determined. Results were reported as percent killing based on luciferase activity in wells with tumor cells alone.

### CD107a degranulation

To assess CD107a degranulation, we plated 1×10^5^ T cells and 5×10^5^ stimulator target cells per well in round-bottom 96-well plates, to a final volume of 200 μL in complete R10 medium, in triplicates. CD107a-PE antibody (BD) was added into each well and incubated at 37°C for 4 hours, along with surface staining for CD8 (Biolegend, San Diego, CA) and CD3 and then analyzed by flow cytometry.

### Flow cytometry

For CAR detection, cells were stained with biotinylated protein L (GenScript, Piscataway, NJ), goat anti-mouse IgG, and anti-human IgG (Jackson ImmunoResearch Laboratories, West Grove, PA), followed by streptavidin-conjugated Allophycocyanin (APC) (BD). Surface expression of EGFR and its mutants was detected by CFP and APC-conjugated cetuximab antibody (Novus Biologicals, Centennial, CO). EGFRvIII expression was detected by anti-EGFRvIII Antibody, clone DH8.3 (Santa Cruz Biotechnology, Dallas, TX). Flow analysis done by LSR Fortessa (BD) and data were analyzed by FlowJo software (BD).

### Animal experiments

All mouse experiments were conducted according to Institutional Animal Care and Use Committee (IACUC)–approved protocols. NSG mice were injected with 2.5×10^5^ U87MG-EGFR/EGFRvIII/Luc+ tumors subcutaneously in 100 μL of PBS on day 0, 7 animals per cohort. Tumor progression was evaluated by luminescence emission on a IVIS Lumin III In Vivo Imaging System (Caliper Life Sciences, Hopkinton, MA) after intraperitoneal D-luciferin injection according to the manufacturer’s instructions (GoldBio). Tumor size was measured by calipers in two dimensions and approximated to volume using the following calculation:

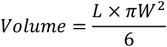

Seven days after tumor implantation, mice were treated with 3×10^6^ CAR T cells intravenously via the tail vein, in 100 μL of PBS. Survival was followed over time until predetermined IACUC-approved endpoints were reached.

### GBM organoids

GBM organoids (GBOs) were established from primary patient tissue, under a University of Pennsylvania Institutional Review Board approved protocol and with patient written informed consent, and co-cultured with CAR T cells as described previously (22, 23). GBOs were fixed and stained after co-culture, using anti-CD3 (BioLegend), anti-cleaved caspase 3 (Cell Signaling Technology, Danvers, MA), anti-EGFR (Thermo Fisher Scientific), anti-EGFRvIII (Cell Signaling Technology), and DAPI (Sigma). To control for tumor heterogeneity, 4 GBOs per condition were used. Mutational data and variant allele fractions (VAF) were obtained from the Center for Personalized Diagnostics at the University of Pennsylvania, as described previously (24).

### Statistical analysis

All in vitro experiments were performed at least in triplicate. GraphPad Prism 6 software (GraphPad Software, San Diego, CA) was used for statistical analyses. Data were presented as mean ± standard deviation. The differences between means were tested by appropriate tests. For the mouse experiments, changes in tumor radiance from baseline at each time point were calculated and compared between groups using t-test or Wilcoxon rank-sum test, as appropriate. Survival determined from the time of T cell injection was analyzed by the Kaplan-Meier method and differences in survival between groups were compared by log-rank Mantel-Cox test.

## Results

### Generation of 806 CARs and Cell lines expressing EGFR mutated proteins

In the present study, we have generated CARs that target EGFR and EGFR mutants by fusing the scFv derived from mAb806 to a 2^nd^ generation CAR construct containing 4-1BB-CD3ζ signaling 806 CAR, the design of which is shown schematically in Fig. 1A. The EGFRvIII-specific 4-1BB-CD3ζ based 2173 CAR used in our clinical trials (NCT02209376 and NCT03726515) was generated for comparative evaluation with 806 CAR. 4-1BB based Cetuximab (C225) and CD19 CARs were used as positive and negative controls. Lentiviral vectors encoding CARs were transduced into a mixture of CD4 and CD8 T cells and surface expression was confirmed by flow cytometry (Fig. 1B). We next turned to generating target-positive tumor cell lines, expressing the mutations EGFR^R108K/G^, EGFR^A289D/T/V^, EGFR^G598V^, and EGFRvIII, for testing of our CAR constructs (Fig. 1C). In order to more faithfully model the EGFR mutations, which are almost always co-expressed with amplified wtEGFR, we transduced the GBM cell line U87MG and patient-derived glioma stem cell line GSC5077 (21), both of which express low levels of wtEGFR, with a lentiviral vector encoding wtEGFR (Fig.1D) (resultant lines referred to as U87MG-EGFR and GSC5077-EGFR), as well as K562 chronic myelogenous leukemia (CML) cells that lack endogenous expression of EGFR, with wtEGFR (Fig. 1E). U87MG-EGFR, GSC5077-EGFR, and K562 cells were also transduced with EGFRvIII lentivirus and expression was then analyzed by an EGFRvIII-specific antibody (Fig.1F). A lentiviral vector co-expressing CFP and the targeted EGFR extracellular mutants (Fig.1D) was transduced into U87MG-EGFR, GSC5077-EGFR, and K562 cells. The resulting CFP positive cells were sorted by fluorescence-activated cell sorting to obtain a positively transduced cell population (Fig. 1G).

**Figure 1.**
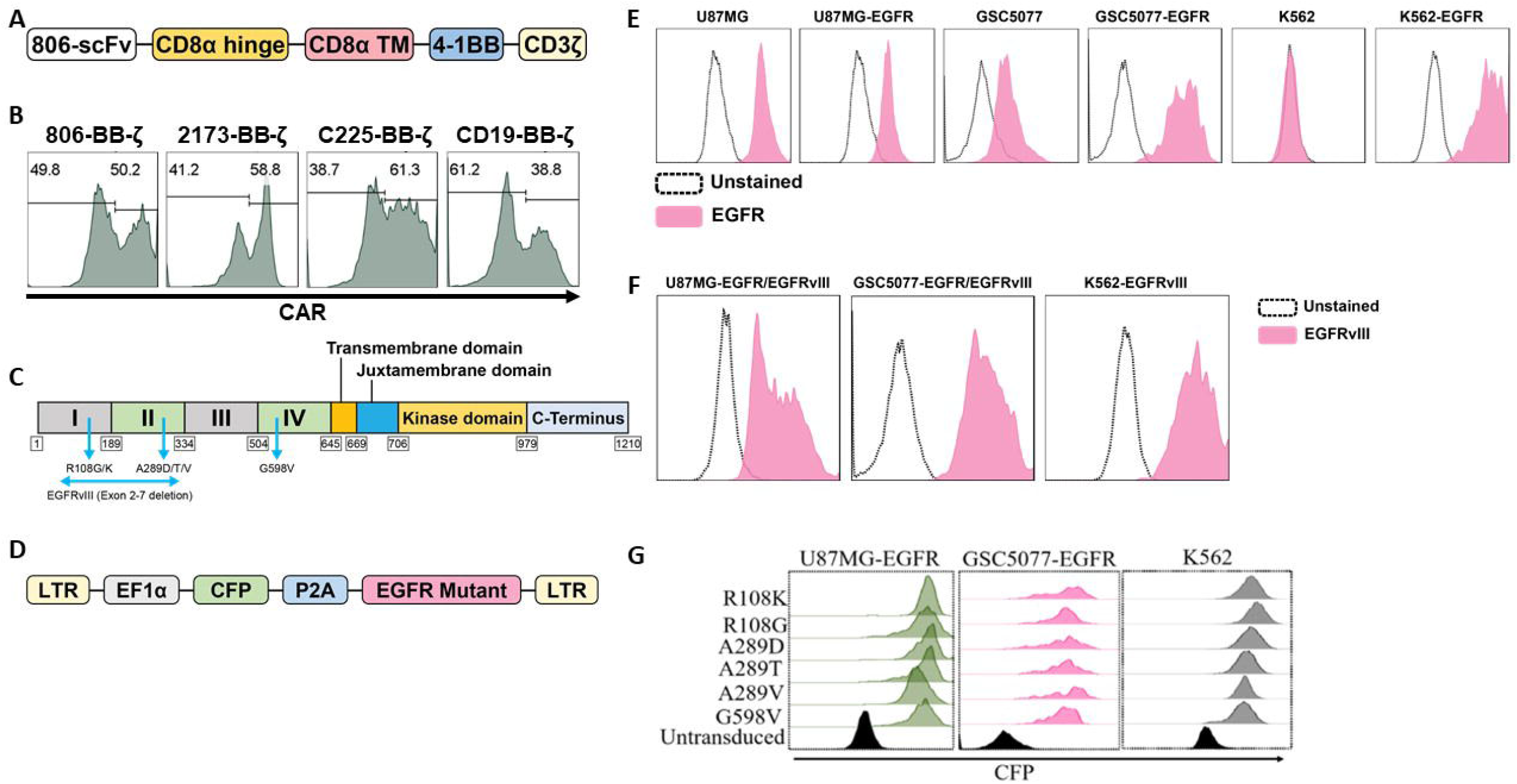
Construction and expression of 806 CAR and EGFR mutant cell lines. (A) Schematic diagram of vector map of 806 CAR containing the 4-1BB co-stimulatory domain. (B) CAR surface expression in primary human CD4^+^ and CD8^+^ T cells. Human T cells were simulated for 24 hours with anti-CD3/anti-CD28 T-cell activating beads and transduced with CAR transgenes and CAR expression was analyzed by flow cytometry using biotinylated goat-anti-mouse (806, C225, and CD19 CARs) and goat-anti human F(ab)2 fragment specific antibodies (2173 CARs) followed by secondary staining with streptavidin-APC. (C) Schematic showing targeted missense mutations in extracellular domain of EGFR, EGFR^R108K/G^, EGFR^A289D/T/V^, EGFR^G598V^ and splice variant EGFRvIII. (D) Schematic of lentiviral vector co-expressing CFP and wtEGFR or EGFR mutant. (E) Flow based analysis of endogenous and ectopically expressed EGFR in U87MG, GSC5077, and K562 cell lines using the cetuximab antibody. (F) U87MG, U87MG-EGFR and GSC5077-EGFR expression of EGFRvIII. (G) U87MG-EGFR, GSC5077-EGFR, and K562 cell lines were transduced with a lentiviral vector co-expressing CFP and indicated EGFR missense mutations and sorted by CFP expression using fluorescent activated cell sorting.

### *In vitro* characterization of 806 CAR T cells

To determine the specificity of the 806 and 2173 CARs for overexpressed wtEGFR, EGFRvIII, and the EGFR-ECD mutants, 2173 and 806 EGFR BB-ζ CAR T cells were co-cultured with U87MG-EGFR and GSC5077-EGFR cell lines expressing EGFRvIII and extracellular mutants EGFR^R108K/G^, EGFR^A289D/T/V^, and EGFR^G598V^, in 24-hour bioluminescence-luciferase based killing assays (Fig. 2A-B). While 2173 CAR T cells demonstrated specificity for EGFRvIII alone, 806 CAR T cells efficiently lysed all targets and exhibited similar cytolytic potential as C225 CAR T (Fig. 2A-B). Notably, 806 CAR T cells were able to kill U87MG cells, despite expressing only low levels of wtEGFR, at an equal level when compared to overexpressed wtEGFR and EGFRvIII. Since U87MG-EGFR mutants expressed endogenous and ectopic EGFR, we could not distinguish if the 806 scFv binding specificity was restricted to the mutant or wtEGFR. To test the exclusive specificity to the mutants, we co-cultured 806 and 2173 CAR T cells with the CML cell line K562, transduced to express wtEGFR, EGFRvIII, or EGFR-mutants, as K562 does not have any endogenous EGFR (Fig. 2B). 806 CAR T cells did not lyse untransduced K562 cells, confirming the lack of EGFR on the parental line. The 806 CAR T cells selectively targeted K562 cells expressing EGFR, EGFRvIII, or EGFR-ECD mutants and demonstrated similar efficacy as C225 CAR T cells. 2173 CAR T cells lysed K562-EGFRvIII cells but did not show any activity against either wtEGFR or the ECD mutants, as expected (Fig. 2B). T cell activation was assessed by induction of surface CD107a expression after co-culture of CAR T cells with target-expressing cells (Fig. 2C). Antigen-specific effector cytokine production was assessed by co-culturing K562 target cells transduced with EGFR and its variants with CAR T cells. The resulting supernatants were analyzed for IFN-γ, TNF-α, and IL2 production (Fig. 2D). Untransduced K562 and Nalm6 cells were used as negative controls. 806 and C225 CAR T cells produced similar levels of CD107 degranulation (Fig. 2C) and IFN-γ, TNF-α, and IL2 (Fig. 2D) in response to EGFRvIII, EGFR-ECD mutants, and EGFR overexpressing cells, while 2173 CAR T cells responded to EGFRvIII alone.

**Figure 2.**
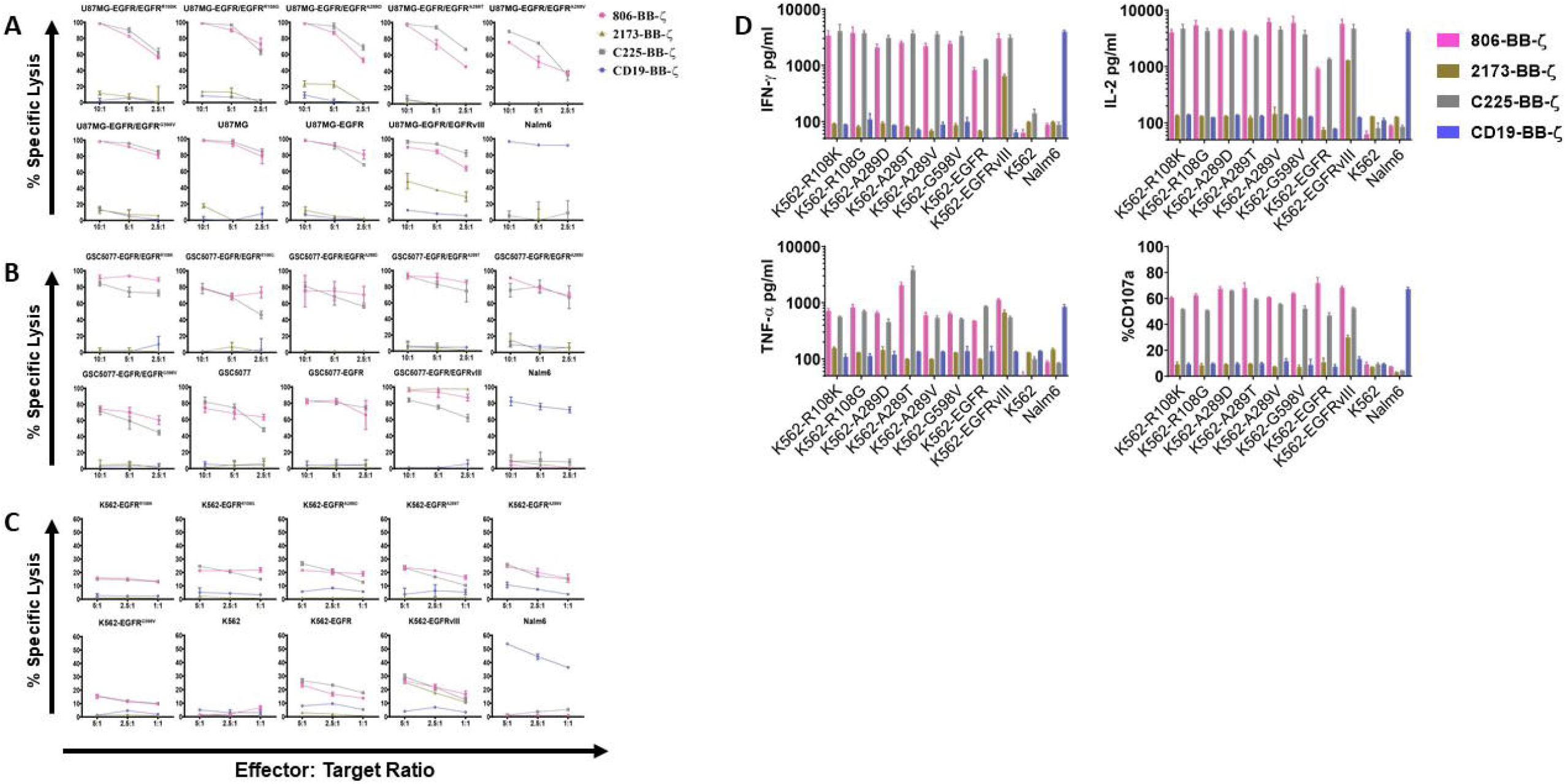
*in vitro* characterization of 806 EGFR CAR T cells. Antigen specific cytolytic activity of 806 and 2173 CAR T cells against cell lines expressing EGFR and its variants. (A) U87MG-EGFR and GSC5077-EGFR cell lines expressing EGFRvIII, EGFR^R108K/G^, EGFR^A289D/T/V^, and EGFR^G598V^ mutant variants were stably transduced with Click Beetle Green (CBG) and co-cultured with CAR T cells at indicated effector to target ratios for 24 hours. One representative experiment from 3 normal donors is shown. Samples were performed in triplicates in 3 replicative experiments. C225-BB-ζ CAR and CD19-BB-ζ CAR were used as positive and negative controls, respectively. (B) Antigen specific cytolytic activity of 806 and 2173 CAR T cells in EGFR and its variants expressed in K562 cells in a 4-hour chromium release assay at indicated effector to target ratios. (C) K562 cells expressing wtEGFR, EGFRvIII, or EGFR-mutants were co-cultured with 806 CART cells for 48 hours. IFN-γ, TNF-α, and IL2 secretion was measured in the supernatant by ELISA. Bar charts represent results from single experiment and values represent the average ± SD of triplicates. (D) CD107a upregulation on CAR T cells stimulated with K562 cells expressing wtEGFR, EGFRvIII, or EGFR-mutants for 4 hours. The percentage of CD107a expression was quantified on CD3 cells (values represent the average of ± SD of 2 repeated experiments).

### 806 CAR T cells exhibit low or no affinity for EGFR expressed on primary astrocytes and keratinocytes

Having confirmed the function of the 806 CARs, we next sought to compare the reactivity of 806 and 2173 CAR T cells in response to endogenous levels of EGFR in normal cells, *in vitro*. We cultured primary human keratinocytes and astrocytes, as those cell types express wtEGFR (Fig. 3A) and used them to stimulate CAR T cells. We observed production of IFN-γ by C225 CAR T cells, in response to EGFR presented by either astrocytes or keratinocytes, as well as U87MG-EGFR (Fig. 3B). In contrast, 2173 CAR T cells produced IFN-γ in response to EGFRvIII antigen alone. 806 CAR T cells exhibited low or no cytotoxicity when co-cultured with astrocytes or keratinocytes (Fig. 3C), with corresponding low IFN-γ production (Fig. 3B).

**Figure 3.**
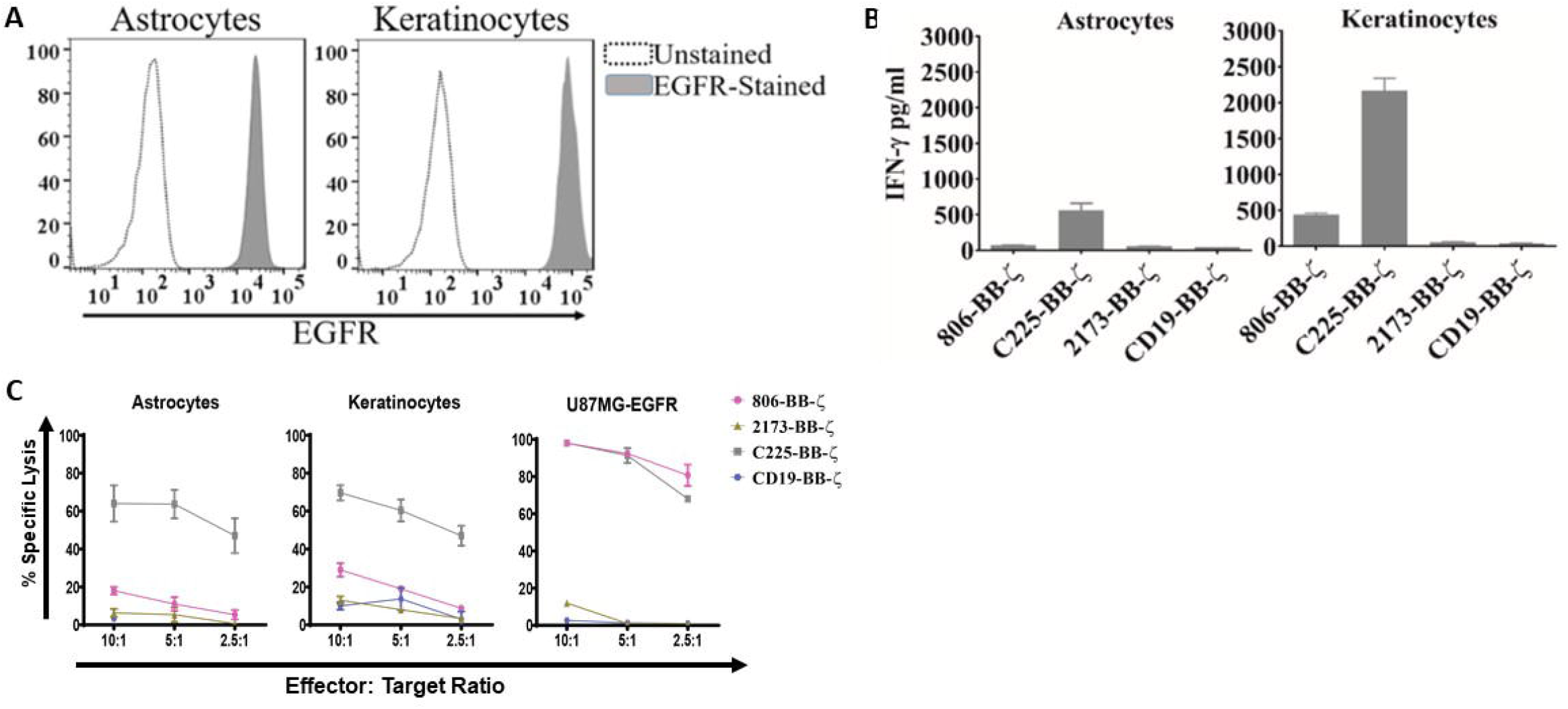
Anti-tumor efficacy of 806 CAR T cells in primary astrocytes and keratinocytes. (A) Surface expression of EGFR assessed by flow cytometry on human primary astrocytes and keratinocytes using EGFR-specific cetuximab antibody. (B) Primary astrocytes and keratinocytes were co-cultured with 806 CAR T cells at indicated ratios in a 4-hour chromium assay and results are representative of a single experiment showing the average ± SD of triplicates. (C) Levels of IFN-γ measured in supernatants by ELISA 24 hours after co-culturing 806 and 2173 CAR T cells with primary astrocytes and keratinocytes at effector to target ratio of 1:1. Results are representative of a single experiment with the average ± SD of triplicates. (D) EGFR expression analysis by quantitative flow cytometry, with quantified EGFR counts on each bar graph.

### Anti-tumor activity of 806 CAR T cells in *in vivo*

Having compared the antigen specific effector function of 806 CAR with 2173CARs, we next sought to confirm its *in vivo* anti-tumor effects, using immunodeficient NSG mice bearing human GBM tumors (Fig. 4A). On Day 0, U87MG-EGFR/EGFRvIII tumors were implanted subcutaneously and on Day 5, tumor engraftment was confirmed by bioluminescence imaging (BLI). On Day 7, a single dose of 3×10^6^ CAR positive T cells were infused intravenously (n = 7 per cohort). Total bioluminescence (Fig. 4B) and individual bioluminescence (Fig. 4C) was assessed in 806 and 2173 CAR T cell treated groups. Animals in the negative control cohort, receiving CD19 CAR T cells, demonstrated rapid tumor growth, with all mice reaching a predetermined humane experimental endpoint by 42 days after initial tumor engraftment. To be noted, all mice in the CD19, 2173, and 806 groups reached experimental endpoint by day 42, 63, and 91 days, respectively (Fig. 4A).

**Figure 4.**
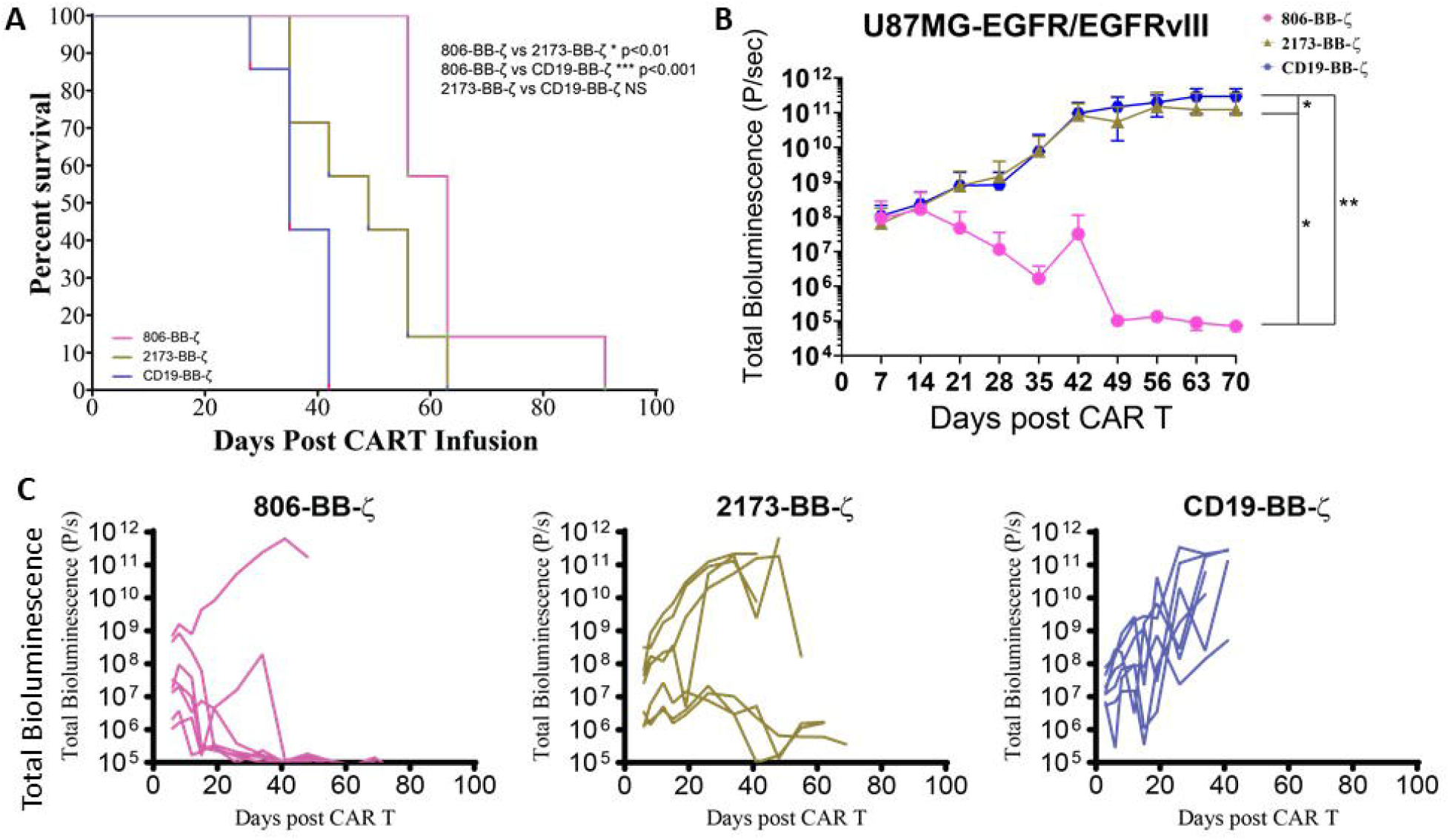
In vivo anti-tumor effect of 806 CAR T cells in NSG mice bearing U87MG-EGFR/EGFRvIII^+^ xenografts. Seven days after 250,000 U87MG-EGFR/EGFRvIII cells were subcutaneously implanted into mice, 3×10^6^ T cells were injected intravenously with indicated CAR constructs. (A) Survival based on time to endpoint was plotted using a Kaplan-Meier curve and statistically significant differences between CAR groups were determined using log-rank Mantel-Cox test. Tumor burden was assessed by bioluminescent imaging. Bars indicate means ± SD (n = 7 mice per group). Tumor burden was quantified as total flux (B) and in individual mice (C) in units of photons/second. Bars indicate means± SD (n = 7 mice). P = photons.

### High fidelity GBM organoids demonstrate cross-reactivity of 806 CAR

Given the ability of the 806 CAR to target EGFR alterations beyond EGFRvIII, we turned to patient-derived GBM organoids (GBOs) to demonstrate activity in a heterogenous model previously characterized to be of high fidelity to human tumors (22, 23). GBOs retain the originating tumor heterogeneity to a high degree out beyond 12 weeks of culturing and maintain expression of endogenous EGFR and its alterations, providing a valuable model platform for testing therapies aimed at addressing tumor escape. The GBOs selected for co-culture experiments contained multiple EGFR mutations (Fig. 5A). GBO 9057 had EGFR copy number gain, EGFRvIII, and two missense mutations, EGFR^G598V^ and EGFR^C595Y^. The missense mutation was found to have a VAF of 24%, while EGFRvIII was identified in less than 10% of the reads, based on next generation sequencing (NGS). GBO 9066 had EGFR copy number gain, EGFR^A289V^, and EGFR^G598V^. Both EGFR^A289V^ and EGFR^G598V^ had a VAF of less than 15%, making determination of co-occurrence impossible through NGS.

**Figure 5.**
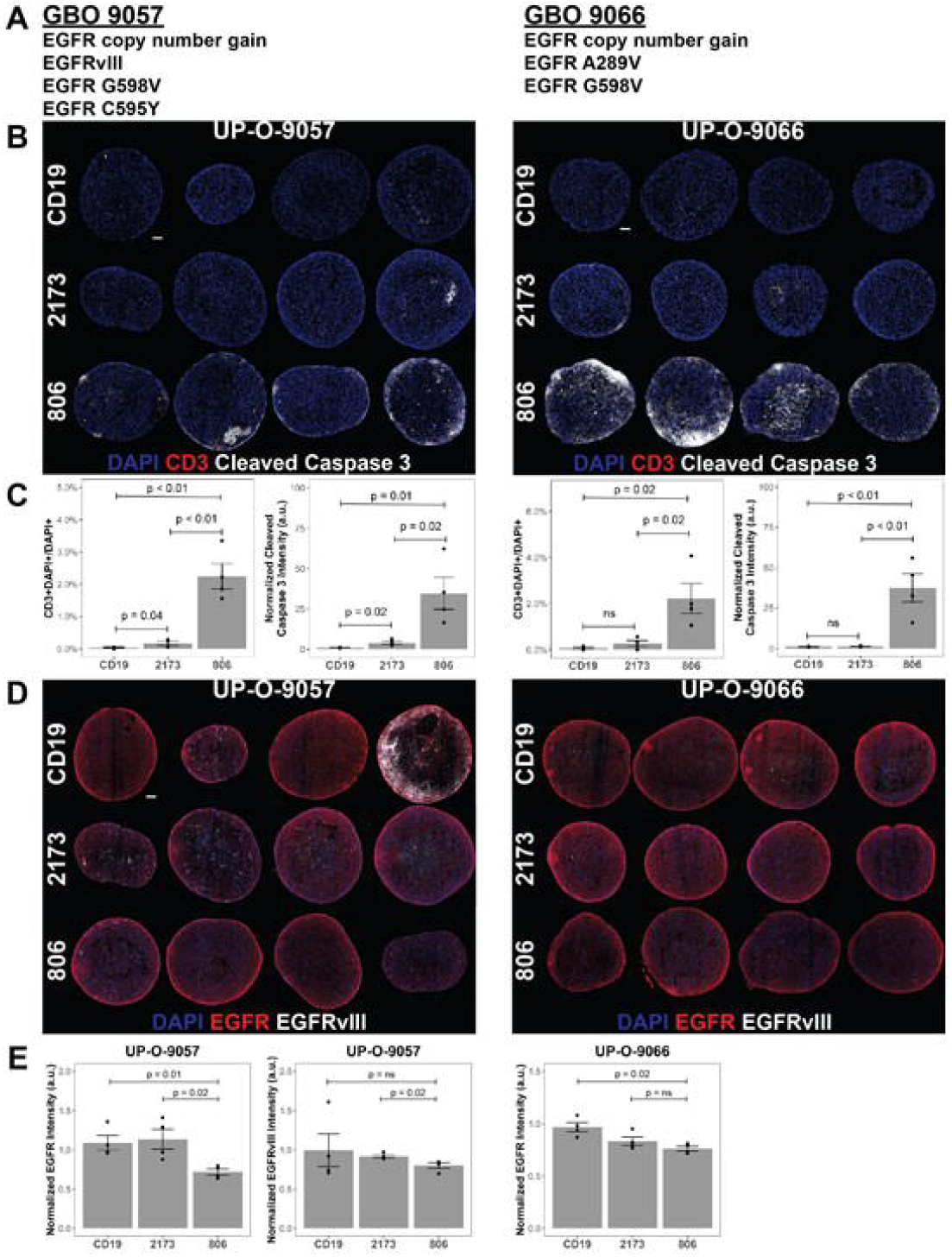
806 CAR T activity in heterogenous GBOs highlights cross-reactivity of 806 binder against oncogenic EGFRs. CAR T co-culture with GBOs was used to demonstrate anti-EGFR activity. (A) EGFR alterations identified in each GBO line. (B) Immunofluorescence images of CAR T cells engrafted GBOs, for 4 organoids per condition, 9057 (left) and 9066 (right). Blue=DAPI; Red=CD3^+^; White=Cleaved caspase 3^+^ (CC3^+^), scale bar = 100 µm. (C) Quantification of CD3^+^ cells (left) and CC3^+^ cells (right) showing anti-tumor activity from the 806 CAR T cells. (D) Immunofluorescence images of CAR T cell targets in GBOs, for 9057 (left) and 9066 (right). Blue=DAPI; Red=EGFR^+^; White=EGFRvIII^+^, scale bar = 100 µm. (E) Quantification of EGFR^+^ (left) and EGFRvIII^+^ signals (right) showing anti-tumor activity from the 806 CAR T cells. Error bars are ± standard error.

GBOs were co-cultured with 806, 2173, and CD19 CAR T cells, at a 1:10 E:T ratio, for 72 hours before fixation and evaluation. CAR T cell infiltration, as quantified by CD3 staining, was more significant in the 806 CAR T cell population than either the 2173 or CD19 CAR T cell population (Fig. 5B). Cleaved caspase 3 (CC3) was used as a measure of cell death and anti-tumor activity. As with the CD3^+^ cell infiltration, the 806 CAR co-culture resulted in higher CC3 levels than either the 2173 or CD19 CAR co-cultures (Fig. 5C). These results highlighted the broad cross-reactivity of the 806 CAR in a heterogeneous, high-fidelity GBM model. wtEGFR staining in both GBO lines provided additional evidence of the cross-reactive nature of the 806 CAR (Fig. 5D). Staining intensity, normalized to CD19 CAR-treated GBOs, showed consistent decreases in 806 CAR-treated GBOs, to a greater degree than the 2173 CAR-treated GBOs (Fig. 5E).

## Discussion

We have shown broad cross-reactivity of 806 CAR T cells to EGFR mutant proteins resulting in enhanced anti-GBM tumor killing, along with a low on-target, off-tumor effect against both astrocytes and keratinocytes that express wildtype EGFR. Importantly, 806 CAR T cells are able to more effectively control tumor growth in a wtEGFR/EGFRvIII model. 806 CAR T cells also demonstrate greater killing in GBOs with heterogeneity of endogenous EGFR and EGFR mutants, confirming its potential to more effectively treat GBM tumors by limiting the impact of tumor escape due to antigen loss.

With regard to the CAR T trial in recurrent GBM (12), the demonstrated tumor recurrence was likely due to the exclusive specificity of the scFv employed in the trial. The 2173 construct was chosen for its selective binding to a novel glycine residue formed at the exons 2-7 deletion in EGFRvIII and for a lack of cross-reactivity to wtEGFR (20). However, the binding affinity and target repertoire were of secondary importance. Given the co-occurrence of amplified wtEGFR with EGFRvIII and most ECD missense mutations (11), there is a clinically-relevant rationale for targeting multiple EGFR alterations in the GBM population (25). Dual targeting of EGFR and EGFRvIII by CAR T and NK cells has been demonstrated in recent studies using scFvs specific for both antigens (26–29). Our work expands on that, as 806 CAR T cells were able to lyse GBM (U87MG, GSC5077) and non-GBM (K562) cell lines modified to express not only wtEGFR and EGFRvIII, but also EGFR extracellular mutations. In comparison, 2173 CAR T cells exhibited specificity for EGFRvIII alone (12, 20).

GBM tumors are significantly heterogeneous, both intratumorally (30, 31) and intertumorally (6). Intratumorally, there are mixed cytological subtypes, exhibiting regional differences in gene expression, key genetic mutations, and chromosomal alterations. This polyclonal nature contributes to therapeutic resistance and tumor escape (32). To address intratumoral heterogeneity, relevant targeted therapies would ideally be able to target larger tumor cell populations within the entire tumor bulk. Given the co-occurrence of wtEGFR amplification seen with EGFR mutations and splice variants (24), the cross-reactive EGFR-targeting 806 scFv should provide greater tumor cell coverage, resulting in better tumor control. The potential for broader tumor control was demonstrated through the high-fidelity, heterogeneous GBO model (22). While the VAFs associated with the originating tumors of the GBOS allow for hypothesizing of independent EGFR mutant tumor populations, one caveat is that the NGS methods used do not allow for concrete determination of subpopulations. The data was subject to bias from tumor viability and number of reads of the sample. Additionally, the heterogeneity of the EGFR variants on amplified alleles is complicated by the mechanisms of amplification of EGFR. GBMs frequently harbor double minutes, extrachromosomal sequences of DNA that are acentric and lead to asymmetric distribution to daughter cells (33). This causes increased cell-to-cell heterogeneity of EGFR alterations in GBM.

Intertumoral variation, from patient to patient, reduces the applicable population for targeted therapies. However, there are gene families frequently found altered across GBM (6). In particular, EGFR amplification is found in up to 60% of GBMs. Concurrently with amplification, 30–40% of GBM tumors express the constitutively active mutant variant, EGFRvIII (34). Combined with the intratumoral expression of EGFR variants, these data suggest that targeting the EGFR family of tumor-specific alterations may successfully address both inter- and intratumoral heterogeneity.

Several EGFRvIII targeted agents are currently in development or in clinical trials for the treatment of GBM. Though the preclinical data from experimental studies evaluating these therapies have been promising, their efficacy in the clinic has yet to be conclusively demonstrated (35–37). In a vaccination approach to target the EGFRvIII in GBM patients, a phase III trial for newly diagnosed glioblastoma failed to show overall efficacy despite 60–80% of recurrent tumors showing complete loss of EGFRvIII positive cells (38). Additional trials targeting EGFRvIII demonstrated similar loss of EGFRvIII concurrent with tumor recurrence (12, 39, 40). Similarly, the EGFRvIII-targeting CAR T trial illustrated the continued presence of EGFR amplification and oncogenic EGFR ECD missense mutations despite EGFRvIII antigen loss in post-treatment tumor specimens (12). These results confirm the need to target multiple EGFR alterations simultaneously.

Despite preclinical efficacy, the success of wtEGFR targeting mAbs cetuximab and panitumumab have been associated with on-target, off-tumor toxicity in other tumor types, due to their significant binding to EGFR expressed on normal tissues (41, 42). Their clinical activity in GBM has yet to be successfully demonstrated in large-scale studies. Co-culture of 806 CAR T cells with basal physiologic EGFR-expressing normal tissue cell lines did not lead to significant cell killing in our work. Previous work has suggested the 806 epitope is exposed on both mutated EGFR (EGFRvIII, EGFR^R108G/K^, EGFR^A289D/T/V^) as well as amplified wtEGFR found on tumors, but not accessible on wtEGFR found on normal tissue (43). The wtEGFR differences have been proposed to be due to different post-translational mannose modifications and kinetics of EGFR trafficking in tumors compared to normal tissue (44). Multiple clinical trials with humanized mAb 806 conjugated to a microtubule inhibitor (ABT-414) have demonstrated only low levels of cutaneous toxicity (45–47). The therapeutic window of CAR T cells for tumor-associated antigens relies on the quantitative difference between antigen-overexpressing tumor and antigen-low normal tissue. Preclinical studies targeting EGFR and erbB2 with affinity-lowered CAR T cells have demonstrated potent antitumor effects against high antigen density while sparing low antigen density normal tissue (48–50). The demonstrated cross-reactivity of 806 CAR T cells for EGFR alterations, including amplified wtEGFR, EGFRvIII, and ECD missense mutations, suggests that 806 CAR T cells may be a more efficacious therapeutic strategy to achieve tumor control and prevent tumor escape via target antigen loss.

## Author Contributions

RT and ZB conceived and carried out the experiments, with contributions from LZ, JVZ, and DZ for specific assays. MM, GM, HS, and DO supervised the project. RT and ZB wrote the manuscript and all authors contributed to the final version of the manuscript.

## Acknowledgments

The authors thank the Human Immunology Core at the University of Pennsylvania for providing leukocytes for the described work, the Stem Cell and Xenograft Core at the University of Pennsylvania for assistance with the animal work, and the Small Animal Imaging Facility at the University of Pennsylvania for the bioluminescence imaging.

